# Distinct roles of parvalbumin and somatostatin interneurons in the synchronization of spike-times in the neocortex

**DOI:** 10.1101/671743

**Authors:** Hyun Jae Jang, Hyowon Chung, James M. Rowland, Blake A. Richards, Michael M. Kohl, Jeehyun Kwag

## Abstract

Synchronization of precise spike-times across multiple neurons carries information about sensory stimuli. Inhibitory interneurons are suggested to promote this synchronization, but it is unclear whether distinct interneuron subtypes provide different contributions. To test this, we examined single-unit recordings from barrel cortex *in vivo* and used optogenetics to determine the contribution of two classes of inhibitory interneurons: parvalbumin (PV)- and somatostatin (SST)-positive interneurons to spike-timing synchronization across cortical layers. We found that PV interneurons preferentially promote the synchronization of spike-times when instantaneous firing-rates are low (<12 Hz), whereas SST interneurons preferentially promote the synchronization of spike-times when instantaneous firing-rates are high (>12 Hz). Furthermore, using a computational model, we demonstrate that these effects can be explained by PV and SST interneurons having preferential contribution to feedforward and feedback inhibition, respectively. Our findings demonstrate that distinct subtypes of inhibitory interneurons have frequency-selective roles in spatio-temporal synchronization of precise spike-times.

## Introduction

Precisely timed spikes that are spatially coordinated or synchronized across multiple neurons with millisecond temporal precision have been shown to encode sensory information about stimuli^1–6^. Information is contained in both the spike times^2,5^ as well as the instantaneous firing-rate (iFR) of precisely timed spike sequences^1,3^, emphasizing the coexistence of temporal and rate codes during sensory information processing^7–9^. Yet, the neural circuit mechanisms supporting the generation of highly synchronized spike sequences across cortical layers remain unknown.

One potential mechanism for spatio-temporal synchronization of precise spike-times is inhibition. Theoretical as well as experimental studies have suggested that inhibition can modulate spatial correlation/synchronization of spike-times between nearby neurons^10–12^ and in neurons across multiple neuronal layers^13,14^. In fact, the latency between excitation and inhibition (E/I latency) has been shown to modulate timing and rate of spike sequences in tandem *in vivo*^7–9^. Thus, E/I latency may have critical role in spatio-temporal synchronization of spike-times. Biologically, differences in E/I latency may be a result of distinct contributions from sensory-evoked feedforward^15–17^ and feedback^18,19^ inhibition. Feedforward inhibition is recruited by afferent inputs that co-activate the inhibition and the neurons being inhibited while feedback inhibition is recruited by activation of the same excitatory neurons that subsequently receive the inhibition. As such, feedback inhibition has a slower onset latency than feedforward inhibition^15,20^. Distinct subpopulations of cortical interneurons, such as parvalbumin (PV)- and somatostatin (SST)-positive inhibitory interneurons, are thought to provide distinct contributions to feedforward and feedback inhibition pathways, with perisomatic-targeting PV interneurons preferentially acting in a feedforward manner on excitatory neurons^21–25^ and dendritic-targeting SST interneurons preferentially acting via feedback pathways to excitatory neurons^22,24,26,27^. Altogether, we are presented with the following picture from the existing literature: inhibition is important for spike-timing synchronization, and it is likely that feedforward and feedback inhibition control spatio-temporal spike-timing synchronization differently, depending on the iFR of inter-synchrony interval. At the same time, PV versus SST interneurons appear more involved in feedforward and feedback inhibition, respectively. Given these considerations, it is important to answer the following questions: (1) Do PV and SST interneurons make distinct contributions to spatio-temporal synchronization of precise spike-times? (2) Are the contributions of PV and SST interneurons to spike-timing synchronization a function of the underlying iFR of the spike sequence? (3) If any differences in the role of PV and SST interneurons in spike-timing synchronization exist, can they be ascribed to their distinct contributions to feedforward and feedback inhibition pathways in the neocortical microcircuit?

Here, we answer these three questions using *in vivo* single-unit recordings across all layers of the primary somatosensory cortex (S1). We find that the whisker-evoked spike-times and their sequences are precisely synchronized between the granular layer (layer 4) and sub-granular layers in subpopulation of neurons (layers 5-6). Using optogenetic perturbations of PV and SST interneurons, we demonstrate that both PV and SST interneurons promote the synchronization of precise spike-times through these pathways, but with distinct contributions depending on the iFR of inter-spike interval (ISI) of the granule layer. Specifically, when the iFR of ISI is low (<12 Hz), PV interneurons are critical for precise spike-timing synchronization. In contrast, when the iFR of ISI is high (>12 Hz), SST interneurons are critical for precise spike-timing synchronization. Furthermore, using a computational model of spike-timing synchronization in a three-layered network with different levels of feedforward and feedback inhibition, we find that these results can be explained by a greater contribution to feedforward inhibition from PV interneurons, and a greater contribution to feedback inhibition from SST interneurons. To our knowledge, our data provide the first ever direct evidence for a role of specialized inhibitory circuit motifs in the neocortex for the spatio-temporal synchronization of precise spike-times. This may be critical to information processing in the neocortex.

## Results

### Synchronization of whisker stimulation-evoked spike-times between granular and sub-granular layers in S1

In order to investigate the synchronization of whisker stimulation-evoked spike-times *in vivo*, we performed single-unit recordings from cortical layers (L) 2/3, 4, 5 and 6 in S1 of anesthetized mice using a 32-channel silicon probe while stimulating whiskers (Fig. 1a; see Methods). Recordings of single-unit activity were assigned to cortical layers using current-source density profiles (Supplementary Fig. 1) and DiI tracks of the silicon probe (Fig. 1b). Based on waveform asymmetry and spike width, we sorted the units into broad-spiking putative excitatory neurons and narrow-spiking putative inhibitory interneurons^28,29^ (Fig. 1c). Only spikes from putative excitatory neurons that showed whisker stimulation-evoked increase in firing rate in the peri-stimulus time histogram (PSTH) were used for further analysis^28^ (Fig. 1d, see Methods).

**Fig. 1.**
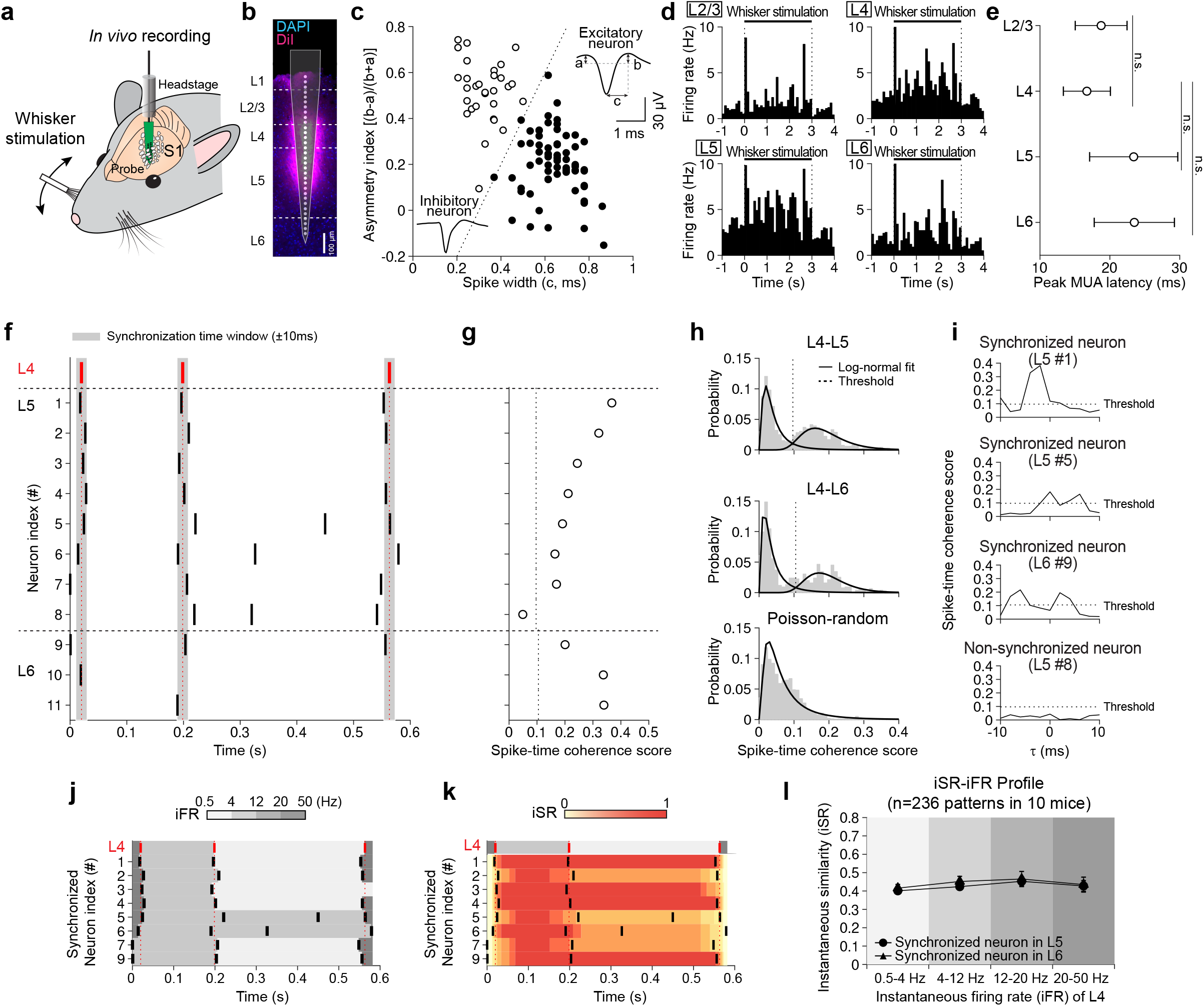
Synchronization of whisker stimulation-evoked spike-times between granular and sub-granular layers in S1 **a** Electrophysiology recordings in S1 during whisker stimulation *in vivo*. **b** Estimated location of the 32-channel silicon probe and contact sites in relation to cortical layers (magenta: Dil-stained probe track, blue: DAPI). **c** Spike waveform-based neuron classification in asymmetry index [(b−a)/(b+a)] versus spike width (c) plot. Dotted line: decision boundary. Inset: initial baseline-to-peak amplitude (a), last baseline-to-peak amplitude (b) and spike width (c) of putative excitatory (filled circles) and inhibitory (empty circles) neurons. **d** Whisker stimulation-evoked changes in firing rate for four representative single units in layers (L) 2/3, 4, 5, and 6. Timepoint 0 denotes whisker stimulation onset. Black bar: whisker stimulation. **e** Peak latency of whisker stimulation-evoked multi-unit activity (MUA) in L2/3, L4, L5, and L6 (n = 4 mice). Data represent mean peak MUA ± SEM. **f** Spike raster plot of putative excitatory neurons in L4, L5, and L6 from one recording session. Dotted red lines indicate the spike-times of four L4 spikes. Gray shade indicates the synchronization time window (±10 ms). **g** Pair-wise spike-time coherence scores of spike-timing sequence. Circles indicate pair-wise coherence scores between a given neuron in L5 or L6 and the L4 neuron indicated in **f**. Vertical dotted line represents the threshold for classifying synchronized and non-synchronized neurons (see **h, i**). **h** Distribution of pair-wise spike-time coherence scores of neuron pairs in L4-L5 (top), L4-L6 (middle) and L4 and spikes generated from a random Poisson process (bottom), fitted with log-normal distribution (solid curve). Threshold: intersection between two log-normal distributions (dotted line). **i** Representative peak spike-time coherence scores of neuron pairs in L4-L5 (top two panels) and L4-L6 (bottom two panels) versus time lag (τ). Neurons with peak coherence scores above threshold (dotted line) are defined “synchronized neuron” or else “non-synchronized neuron”. **j** Representative plot of instantaneous firing-rate (iFR, four bins: 0.5-4, 4-12, 12-20 and 20-50 Hz, gray-scale) of neurons in L4 and L5/L6. **k** Representative plot of instantaneous similarity (iSR, maximum 1, red color-scale, bottom) of neurons in L4 and L5/L6. **l** iSR-iFR profile of synchronized neurons in L5 (circle) and L6 (triangle). All data are mean ± SEM and n represents the number of animals.

To determine the initial receptive layer that responds to whisker stimulation in S1, we analyzed the latency of the peak multi-unit activity (MUA) of all whisker stimulation-responsive neurons in each layer (Fig. 1e, see Methods). The thalamo-recipient granular layer L4 had the earliest peak, followed closely by L2/3, and then after a longer delay sub-granular L5 and L6, similar to what has been observed in other *in vivo* studies^30,31^. This is consistent with the canonical feedforward neocortical microcircuit that has been previously proposed, wherein L4 is the major recipient of primary sensory information from the thalamus^32,33^. However, the peak MUA latencies were heterogeneous across trials, and they were not statistically different (Peak MUA latency of L2/3: 18.75 ± 3.71 ms, L4: 16.71 ± 3.37 ms, L5: 23.41 ± 6.31 ms, L6: 23.5 ± 5.74 ms; n = 4 mice, F_(3, 12)_ = 0.476, *p* = 0.71, one-way ANOVA test; Fig. 1e). This is indicative of non-canonical routes for information flow through cortical layers, for example, through direct connections between the thalamus and L5^34^. However, our investigation focused on the synchronization of whisker stimulation-evoked spike-times between L4 and L5/L6, in-line with the canonical model^32,33^. Neurons in L2/3 were not included in our analyses due to limited statistical power resulting from a small number of detected L2/3 neurons, which may have been caused by the sparsity of L2/3 responses^31,35^.

In order to better understand the synchronization of spike-times between L4 and L5/L6, we performed a pair-wise coherence analysis between spike-timing sequences of pairs of single putative excitatory neurons recorded during each whisker stimulation trial, with each pair comprised of one neuron from L4 and one neuron from L5 or L6 (Fig. 1f). We developed a spike-time coherence score for each pair of L4-L5 and L4-L6 neurons (Fig. 1g). This spike-time coherence score measured the extent to which the L5/L6 neurons in the pair reproduced the spike-timing sequences recorded in the L4 neuron during the whisker stimulation trial, allowing for a synchronization time window of ±10 ms (gray shade, Fig. 1f). That is, this coherence score was the normalized cross-correlation of the two spike trains within the synchronization time window (±10 ms, see Methods) and if during whisker stimulation the L5 and L6 neurons in the pair tended to spike within the synchronization time window, the pair would receive a coherence score close to 1, otherwise the pair would receive a coherence score close to zero (Fig. 1f, g).

When we examined the spike-time coherence scores across all pairs and trials, the distributions were clearly bimodal in both L4-L5 and L4-L6 pairs (Fig. 1h, top and middle; Silverman’s test with unimodal null hypothesis of L4-L5: *p* < 0.001, L4-L6: *p* < 0.05). In contrast, spike-time coherence scores between L4 spikes and spikes generated from a random Poisson process had a unimodal distribution (Fig. 1h, bottom; Silverman’s test with unimodal null hypothesis: *p* = 0.22). Moreover, in a surrogate dataset, bimodality of spike-time coherence scores disappeared when we shuffled ISI (Supplementary Fig. 2a-c, top) or Poisson-randomized spike-times (Supplementary Fig. 2a-c, bottom). This suggests that in the real data, on any given trial, a subset of sub-granular neurons do synchronize L4 spike-timing sequences. We note, though, that on any given trial different sets of neurons were more coherent or less coherent (Supplementary Fig. 3a,d), suggesting that the bimodal distributions do not reflect the presence of two fundamentally distinct neuronal populations in the sub-granular layers consistent with previous *in vivo* observation that different synchronized groups may originate in the same or overlapping neuronal populations^3,6^.

In order to focus the scope of our analysis onto those L5 and L6 neurons that synchronize L4 spike-times on any given trial, we fit a mixture of two log-normal distributions to the data using the Expectation-Maximization algorithm^36^ (see Methods). The data was well fit with this model (Fig. 1h; *r*^2^ of L4-L5 = 0.93, L4-L6 = 0.85), and provided us with an empirically determined threshold (Fig. 1h, vertical dotted line) for distinguishing between those neurons that participated in the synchronization of spike-times from L4 that had spike-time coherence score greater than the threshold and those that did not (Fig. 1h, i; threshold of L5 = 0.10, L6 = 0.11; see Methods). We refer to these neurons as “synchronized neurons” and “nonsynchronized neurons”, but we note that the definition applied to neurons during individual trials, so a neuron could switch its status between “synchronized” and “non-synchronized” across different trials (Supplementary Fig. 3a). In the analyses that follow, we only examine “synchronized neurons”.

Since spike-time and firing rate co-exist and are not completely dissociable in the spike-timing sequences^7–9^, it is important to examine how the synchronization of spike-times depend on the iFR of ISI. Thus, we directly compared the spike-times of synchronized L5 and L6 neurons with that of L4 neurons as a function of the L4 neurons’ iFRs. To measure the similarity between spike-times in pairs of L4-L5 and L4-L6 neurons at any given point in time, and to express that as a function of the iFR, we used a previously developed spike-time similarity score that can be applied to sliding windows over time^37^ (see Methods), and we then compared that to an estimate of the iFR in the L4 neurons. Specifically, at any point in time we estimated the iFR of the L4 neuron in the pair as the inverse of the neuron’s ISI (Fig. 1j). Then, we calculated the spike-time similarity score of the spikes in the L5/L6 neurons from the pair within the synchronization time-window (±10 ms), providing an instantaneous similarity measure (iSR) (Fig. 1k). This measured the extent to which these synchronized L5 or L6 neurons were reproducing the spikes observed in the L4 neuron at each moment in time (see Methods). As a result, for each time-point we had both an estimate of the iFR in the L4 neuron in the pair, and a measure of the spike-time similarity in the L5/L6 neuron in the pair. To simplify our analysis, we grouped the iFR into four different bins: 0.5-4 Hz, 4-12 Hz, 12-20 Hz, and 20-50 Hz. We selected these four bins as they correspond to the range of firing-rates and oscillatory frequencies likely to be observed *in vivo*^38^. We then plotted the iSR metric for each bin of iFRs, for each pair of L4-L5/L6 neurons (iSR-iFR profile, Fig. 1l). We found no evidence of a difference between pairs of L4-L5 and L4-L6 neurons in their iSR-iFR profiles (Fig. 1l; L5: F_(3,526)_ = 1.2, *p* = 0.31, L6: F_(3,314)_ = 0.58, *p* = 0.63, one-way ANOVA test). Moreover, we found no evidence for differences in iSR at different iFR of L4 neurons (Fig. 1l; 0.5-4 Hz: *p* = 0.53, 4-12 Hz: *p* = 0.32, 12-20 Hz: *p* = 0.74, 20-50 Hz: *p* = 0.72; Wilcoxon rank-sum test). A similar trend was observed in L2/3, although the recorded neurons were sparse (Supplementary Fig. 4a-e). Altogether, this data demonstrates that, on any given trial, there are heterogeneous subsets of neurons in the sub-granular layers that spatially synchronize spike-timing sequences between L4 and L5/L6 during whisker stimulation, but in a manner that does not differ between L5 or L6 and which is insensitive to the iFR of L4 neurons.

### Optogenetic activation of PV and SST interneurons gates the synchronization of spike-times in a frequency-selective manner

To investigate the role of inhibitory interneurons in the spatio-temporal synchronization of precise spike-times, we activated PV and SST interneurons via conditionally expressed Channelrhodopsin-2 (ChR2) during whisker stimulation (Fig. 2a)^39^. Immunostaining showed that ChR2-mCherry expressed across all cortical layers in PV-Cre (Fig. 2b, left) or SST-Cre mice (Fig. 2b, right). The ChR2 expression was confirmed by light stimulation-induced changes in the firing rate of putative excitatory neurons and PV or SST interneurons, as shown in the PSTH (Fig. 2c, d). Optical stimulation of ChR2-expressing PV (ChR2-PV) and ChR2-expressing SST (ChR2-SST) interneurons increased their firing rates (Fig. 2c, d, left), which in turn decreased the firing rates of some putative excitatory neurons (Fig. 2c, d, right), confirming successful ChR2 expression. We chose a light intensity that only moderately reduced spiking in putative excitatory neurons (Standard deviation (SD) following activation of ChR2-PV interneurons in PV-Cre mice = 5.93%, SD following activation of ChR2-SST interneurons in SST-Cre mice = 8.05%, Supplementary. Fig 5a-d, see Methods) and confirmed that light could activate ChR2 in sub-granular layers (Supplementary Fig. 6a).

**Fig. 2.**
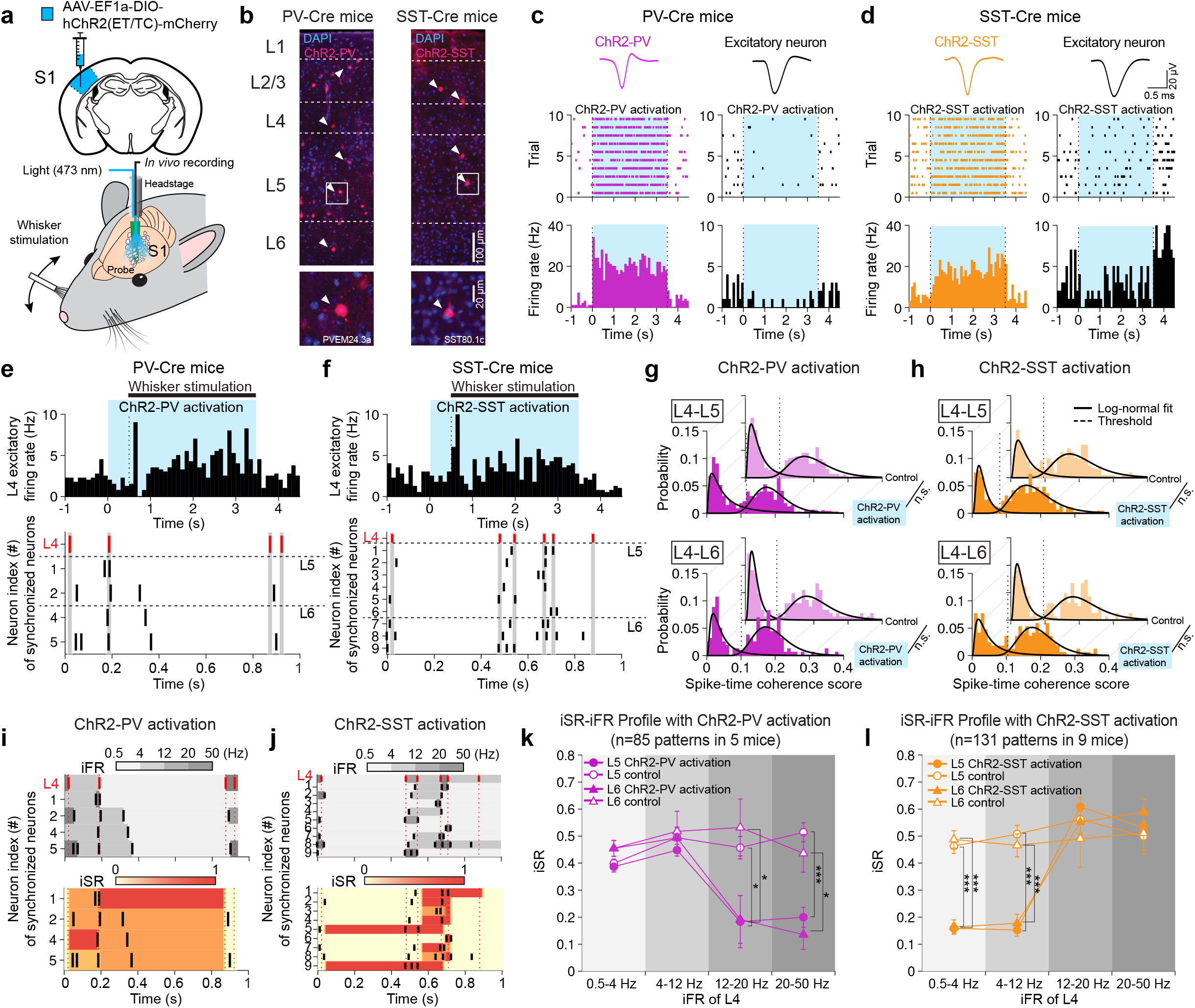
Optogenetic activation of PV and SST interneurons gates the synchronization of spike-times in a frequency-selective manner **a** Top, schematic of the injection of AAV-EF1a-DIO-hChR2(ET/TC)-mCherry (ChR2, blue) into S1. Bottom, electrophysiology recording during whisker stimulation and blue light stimulation (473 nm) in either PV-Cre or SST-Cre mice. **b** ChR2-mCherry-expressing PV (ChR2-PV) interneurons in PV-Cre mice (red, left) and ChR2-mCherry-expressing SST (ChR2-SST) interneurons in SST-Cre mice (red, right) among all cells stained with DAPI (blue). **c** Representative spike waveforms (top), raster plots (middle) and peri-stimulus time histogram (PSTH, bottom) of putative ChR2-PV interneuron (magenta, left) and excitatory neuron (black, right) during blue light stimulation (3.5 s, blue shade) in PV-Cre mice. **d** Same as (**c**) but for putative ChR2-SST interneuron (orange) in SST-Cre mice. **e, f** PSTH of putative L4 excitatory neurons (top) and the raster plots (bottom) of synchronized neurons in L5 and L6 during whisker stimulation (black horizontal bar) with light stimulation (blue shade) in PV-Cre mice (**e**) and SST-Cre mice (**f**). Light stimulation preceded whisker stimulation by 500 ms (dotted lines). Gray shade indicates the synchronization time window (±10 ms). **g, h** Distribution of pair-wise spiketime coherence scores of neuron pairs in L4-L5 (top) and L4-L6 (bottom) with ChR2-PV activation (**g**, magenta) and ChR2-SST activation (**h**, orange) and in control (light magenta/orange), fitted with log-normal distribution (solid curve). Dotted line: threshold between synchronized and non-synchronized neurons. Inset: n.s. *p* > 0.05, Kolmogorov-Smirnov test. **i, j** Representative plot of iFR (four bins: 0.5-4, 4-12, 12-20 and 20-50 Hz, gray-scale, top) and iSR (maximum 1, red color-scale, bottom) of neurons in L4 and L5/L6 during blue light stimulation in PV-Cre mice (**i**) and SST-Cre mice (**j**). Red dotted vertical lines indicate the spike times of the L4 neurons. **k, l** iSR-iFR profiles of synchronized neurons in L5 (circle) and L6 (triangle) during light on (filled) and off (empty) in PV-Cre mice (K, magenta) and in SST-Cre mice (L, orange). All data are mean ± SEM and n represents the number of animals. Inset: \**p* < 0.05, \*\**p* < 0.01, and \*\*\**p* < 0.001, Wilcoxon rank-sum test.

Optical activation of ChR2-PV or ChR2-SST interneurons during whisker stimulation did not interfere with the generation of reliable whisker stimulation-evoked responses in L4 (Fig. 2e, f, top) and L5/L6 (Fig. 2e, f, bottom). As such, we were still able to examine spike-time coherence scores of putative excitatory neurons in L5 and L6 during optical activation of ChR2-PV and ChR2-SST interneurons (Fig. 2g, h). The bimodal distribution of spike-time coherence scores was not significantly affected in pairs of L4-L5 neurons (ChR2-PV in L4-L5: *p* = 0.77, ChR2-SST in L4-L5: *p* = 0.24, Kolmogorov-Smirnov test; Fig. 2g, h, top) and L4-L6 neurons (ChR2-PV in L4-L6: *p* = 0.26, ChR2-SST in L4-L6: *p* = 0.59, Kolmogorov-Smirnov test; Fig. 2g, h, bottom). These data demonstrate that activation of PV and SST interneurons do not have an impact on the switching between synchronous and non-synchronous mode of synchronized neurons (Supplementary Fig. 3b-d). Next, we examined the effect of PV and SST interneuron activation on the spike-timing synchronization could be dependent on the iFR of ISI (Fig. 2i, j). Interestingly, we found very pronounced effects of optical activation of ChR2-PV and ChR2-SST interneurons on the iSR-iFR profiles. Specifically, optical activation of ChR2-PV interneurons led to a large decrease in the spike-time similarity measures of both L5 and L6 neurons at high iFR (Fig. 2k; L5: 12-20 Hz: *p* < 0.05, 20-50 Hz: *p* < 0.001, Wilcoxon rank-sum test; L6: 12-20 Hz: *p* < 0.05, 20-50 Hz: *p* < 0.05, Wilcoxon rank-sum test). In contrast, optical activation of ChR2-SST interneurons led to pronounced decreases in the iSR scores at low iFR (Fig. 2l; L5: 0.5-4 Hz: *p* < 0.001, 4-12 Hz: *p* < 0.001; L6: 0.5-4 Hz: *p* < 0.001, 4-12 Hz: *p* < 0.001, Wilcoxon rank-sum test). The complementary effects of PV and SST interneurons on the iSR-iFR profiles of L5 and L6 putative excitatory neurons suggests that these different classes of inhibitory interneurons act as frequency-selective gates, with PV interneurons gating spike-timing synchronization at high iFR (i.e. acting as a low-pass filter), and SST interneurons gating spike-timing synchronization at low iFR (i.e. acting as a high-pass filter). These results demonstrate that PV and SST interneurons do exert strong, complementary influences on the synchronization of spike-timing sequences from granular to the sub-granular layers.

### Optogenetic inactivation of PV and SST interneurons promotes the synchronization of spike-times in a frequency-selective manner

Our optical activation data demonstrated that PV and SST interneurons can *gate* spike-timing synchronization at specific iFR. We hypothesized that these interneurons might also *promote* spike-timing synchronization in the range of the iFR that they do not gate. In other words, we speculated that if PV interneurons gate spike-times at high iFR, then they may promote spike-timing synchronization at low iFR, and *vice-versa* for SST interneurons. To test this hypothesis, we optogenetically silenced PV and SST interneurons using Archaerhodopsin-3 (Arch)^40^ with 565 nm light (Fig. 3a). Immunostaining showed Arch-EYFP expressed across all cortical layers in PV-Cre or SST-Cre mice (Fig. 3b). Illumination of Arch-expressing PV (Arch-PV) and Arch-expressing SST (Arch-SST) interneurons decreased their firing rates (Fig. 3c, d, left), which in turn increased the firing rates of some putative excitatory neurons (Fig. 3c, d, right), confirming successful Arch expression and activation. We selected a light intensity level that moderately increased the firing rates of putative excitatory neurons (SD following Arch-PV in PV-Cre mice = 4.13%, Arch-SST in SST-Cre mice = 4.64%, Supplementary Fig. 5e-h, see Methods) and we confirmed that light could activate Arch in sub-granular layers (Supplementary Fig. 6b).

Optical silencing of Arch-PV and Arch-SST interneurons did not impact the generation of reliable, whisker-stimulation evoked responses in L4 (Fig. 3e, f, top) and in L5/L6 (Fig. 3e, f, bottom), allowing us to again examine spike-time coherence scores. Interestingly, not as in the case of optical activation of these interneuron classes, we found a moderate change in the spike-time coherence scores between L4-L5 pairs during optical silencing of Arch-PV and Arch-SST interneurons (Arch-PV: *p* < 0.001, Arch-SST: *p* < 0.001, Kolmogorov-Smirnov test), and L4-L6 pairs during optical silencing of Arch-PV and Arch-SST interneurons (Arch-PV: *p* < 0.01, Arch-SST: *p* < 0.01, Kolmogorov-Smirnov test; Fig. 3g, h), implying that PV and SST interneurons might promote the synchronization of precise spike-times while the overall probability of a neuron switching between synchronous and non-synchronous mode was unaffected by optogenetic inactivation (Supplementary Fig. 3b-d). Next, we again examined the iSR-iFR profiles for L5 and L6 neurons in light-on versus light-off conditions (Fig. 3i, j). We found that optical silencing of PV interneurons led to a selective decrease in spike-time similarity scores in both L5 and L6 excitatory neurons at low iFR (Fig. 3k; L5: 0.5-4 Hz: *p* < 0.001, 4-12 Hz: *p* < 0.05; L6: 0.5-4 Hz: *p* < 0.001, 4-12 Hz: *p* < 0.001, Wilcoxon rank-sum test). In contrast, optical silencing of SST interneurons selectively decreased spike-time similarity scores at high iFR (Fig. 3l; L5: 12-20 Hz: *p* < 0.001, 20-50 Hz: *p* < 0.001, Wilcoxon rank-sum test; L6: 12-20 Hz: *p* < 0.05, 20-50 Hz: *p* < 0.01, Wilcoxon rank-sum test). These findings supported our hypothesis, demonstrating that PV and SST interneurons not only gate synchronization of spike-timing sequences, but also promote it, though at different and complementary iFR. Specifically, PV interneurons gate spike-timing synchronization at high-firing rates, and promote it at low-firing rates, whereas SST interneurons gate spike-timing synchronization at low firing rates and promote it at high firing rates.

**Fig. 3.**
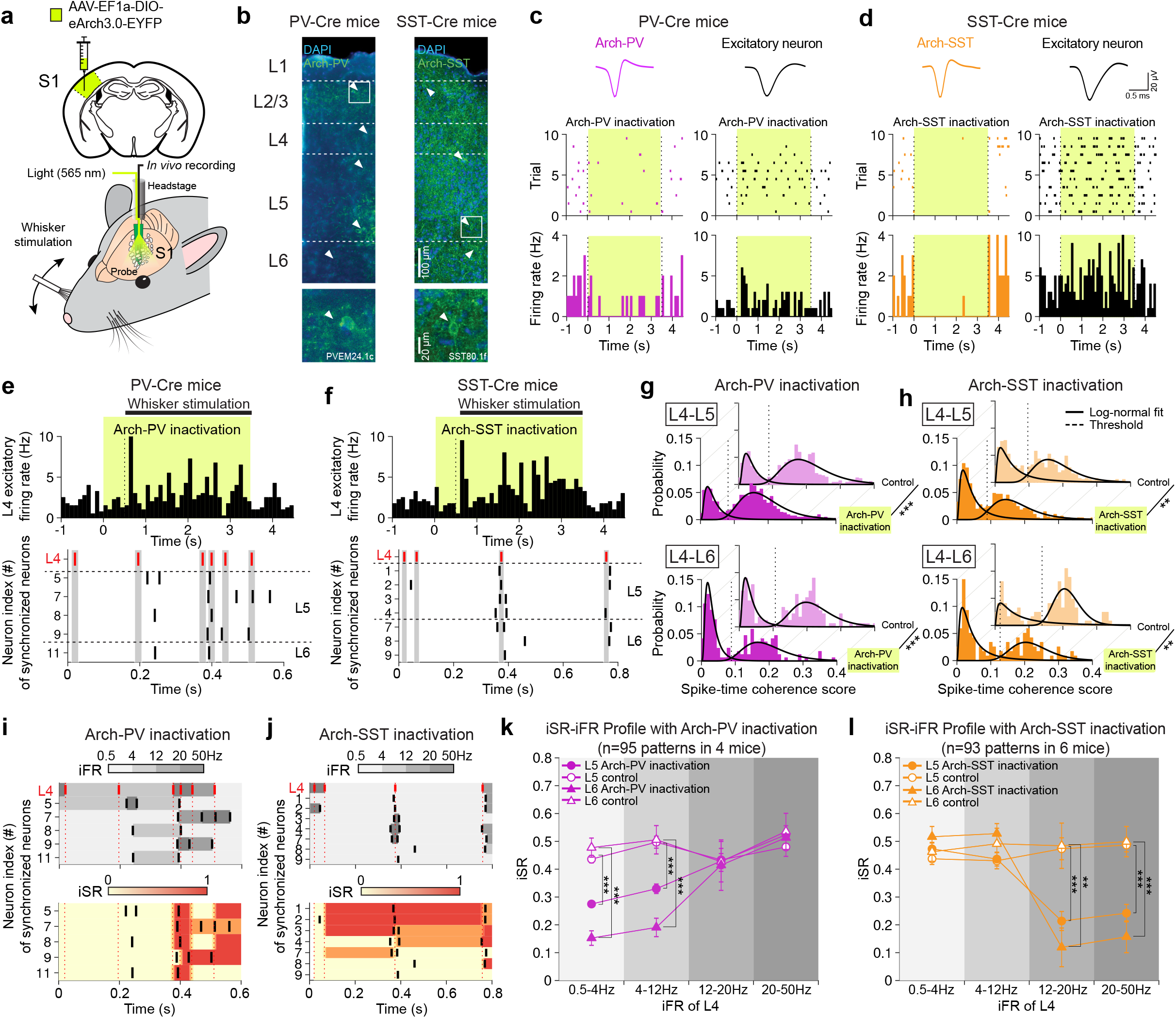
Optogenetic inactivation of PV and SST interneurons promotes the synchronization of spike-times in a frequency-selective manner. **a** Top, schematic of the injection of AAV-EF1a-DIO-eArch3.0-EYFP (Arch, green) into S1. Bottom, electrophysiology recording during whisker stimulation and green light stimulation (565 nm) in either PV-Cre or SST-Cre mice. **b** Arch-EYFP-expressing PV interneurons (Arch-PV) in PV-Cre mice (green, left) and Arch-EYFP-expressing SST interneurons (Arch-SST) in SST-Cre mice (green, right) in EYFP among all cells stained with DAPI (blue). **c** Representative spike waveforms (top), raster plots (middle) and PSTH (bottom) of putative Arch-PV interneuron (magenta, left) and excitatory neuron (black, right) during green light stimulation (3.5 s, green shade) in PV-Cre mice. **d** Same as (**c**) but for putative Arch-SST interneuron (orange) in SST-Cre mice. **e, f** PSTH of putative L4 excitatory neurons (top) and the raster plots (bottom) of synchronized neurons in L5 and L6 during whisker stimulation (black horizontal bar) with light stimulation (green shade) in PV-Cre mice (**e**) and SST-Cre mice (**f**). Light stimulation preceded whisker stimulation by 500 ms (dotted lines). Gray shade indicates the synchronization time window (±10 ms). **g, h** Distribution of pair-wise spike-time coherence scores of neuron pairs in L4-L5 (top) and L4-L6 (bottom) with Arch-PV inactivation (G, magenta) and Arch-SST inactivation (H, orange) and in control (light magenta/orange), fitted with log-normal distribution (solid curve). Dotted line: threshold between synchronized and non-synchronized neurons. Inset: \*\**p* < 0.01 and \*\*\**p* < 0.001, Kolmogorov-Smirnov test. **i, j** Representative plot of iFR (four bins: 0.5-4, 4-12, 12-20 and 20-50 Hz, gray-scale, top) and iSR (maximum 1, red color-scale, bottom) of neurons in L4 and L5/L6 during green light stimulation in PV-Cre mice (**i**) and SST-Cre mice (**j**). Red dotted vertical lines indicate the spike times of the L4 neurons. **k, l** iSR-iFR profiles of synchronized neurons in L5 (circle) and L6 (triangle) during light on (filled) and off (empty) in PV-Cre mice (**k**, magenta) and in SST-Cre mice (**l**, orange). All data are mean ± SEM and n represents the number of animals. Inset: \*\**p* < 0.01 and \*\*\**p* < 0.001, Wilcoxon rank-sum test.

### PV and SST interneurons preferentially recruit feedforward and feedback inhibition in frequency-selective spike-timing synchronization

One hypothesis as to why PV and SST interneurons have these distinct, complementary effects on spike-timing synchronization is that they contribute differentially to feedforward versus feedback inhibition pathways^21,22,24^. However, there is as of yet no experimental technique that can selectively activate or inactivate feedforward and feedback inhibition pathways *in vivo*. Therefore, to address this question, we built a three-layer spiking excitatory neural network model of the sort that has previously been used in studies on the synchronization of precise spike-times^13,41,42^ (see Methods for more details on the computational model). With this model, we could directly implement various inhibitory neural circuit motifs, e.g. motifs with only feedforward inhibition (Fig. 4a, top), only feedback inhibition (Fig. 4a, middle), or some mixture of the two (Fig. 4a, bottom). Furthermore, we could measure iSR-iFR profiles in our *in silico* model, and directly compare them to our *in vivo* data under various conditions of optogenetic perturbation. To calculate iSR-iFR profiles for the *in silico* model, we randomly selected synchronized spike-timing sequences recorded from putative excitatory neurons in L4 during whisker stimulation. We fed these sequences as inputs to the spiking neural network model, and examined the spike-timing sequences that were synchronized to the final layer of the model (Fig. 4b). We then calculated the synthetic iSR-iFR profiles by treating the input layer (L_input_) as the functional equivalent to L4 in S1, and measuring the spike-time similarity scores for neurons in the output layer of the model (L3), treating them as the functional equivalent to sub-granular neurons. Thus, for every pair of L_input_-L3 neurons, we could calculate an iSR-iFR profile, much as we did for L4 and L5/L6 neurons in S1 (Fig. 4c). Intriguingly, we found that in the model with purely feedforward inhibition motifs, precise spike-times only synchronized at low iFRs, as we had observed during optical activation of ChR2-PV interneurons and optical silencing of Arch-SST interneurons (Fig. 4d, top). Treating the *in silico* and *in vivo* iSR-iFR profiles as 4D vectors, we measured the cosine similarity between them, and found that the iSR-iFR profiles in the feedforward inhibition only model was similar to the optical activation of ChR2-PV interneurons and optical silencing of Arch-SST interneurons (Fig. 4e, top). In contrast, in the purely feedback inhibition model, we found that spike-times only synchronized at high iFRs, as we had observed during optical activation of ChR2-SST interneurons and optical silencing of Arch-PV interneurons (Fig. 4d, middle). As expected based on this data, the iSR-iFR profile for the purely feedback inhibition model showed greater cosine similarity to the optical silencing of Arch-PV interneurons and optical activation of ChR2-SST interneurons (Fig. 4e, middle). Notably, in order to obtain a flat iSR-iFR profile, as we had observed in the absence of optical perturbations (Fig. 1l), we found that a mixture of feedforward and feedback inhibition was required. Specifically, a 7:3 mixture of feedforward to feedback inhibition provided the greatest similarity to the control *in vivo* data (Fig. 4d, e, bottom). Altogether, these results suggest that there is a balance between feedforward and feedback inhibition in S1, and our optical manipulations altered spike-timing synchronization by breaking this balance. Specifically, optical activation of ChR2-PV interneurons or optical silencing of Arch-SST interneurons may have promoted feedforward inhibition over feedback inhibition, while optical silencing of Arch-PV interneurons or optical activation of ChR2-SST interneurons may have promoted feedback inhibition over feedforward inhibition (Fig. 4f).

**Fig. 4.**
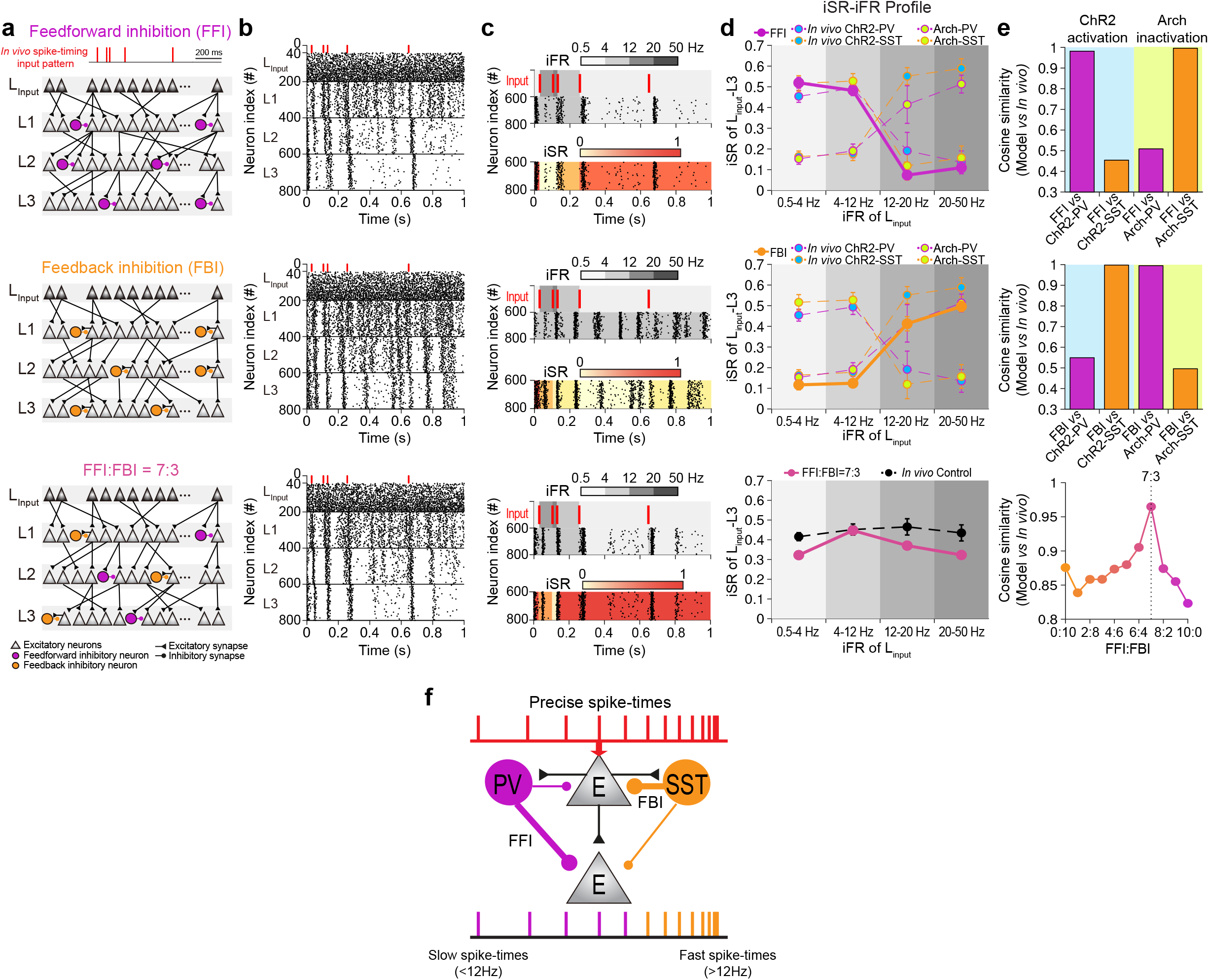
PV and SST interneurons preferentially recruit feedforward and feedback inhibition for frequency-selective spike-timing synchronization. **a** A schematic of a three-layer network model consisting of 200 single-compartment Hodgkin-Huxley excitatory neuron models (triangles) and 50 inhibitory interneuron models (circles) that provide either feedforward inhibition (FFI, magenta, top), feedback inhibition (FBI, orange, middle) and FFI and FBI at 7:3 ratio (pink, bottom). **b** Spike raster plot of each model in response to spike-timing sequence recorded in L4 *in vivo* (red). Dots indicate spikes. A subset of neurons (40 neurons) in the input layer (L_input_) received the input (red dots) while the rest spiked spontaneously (black dots) reflecting sparseness. **c** Representatives of iFR (four bins: 0.5-4, 4-12, 12-20 and 20-50 Hz, gray color-scale, top) and iSR (maximum 1, red color-scale, bottom) of neurons in L_input_-L3 pair in each model. Red in L_input_ are the input spike-timing sequences. Black dots are spikes in L3. **d** iSR-iFR profiles of network model with FFI (magenta circles/line, top), FBI (orange circles/line, middle) and with FFI and FBI at 7:3 (pink circles/line, bottom). *In vivo*-recorded iSR-iFR profiles following ChR2-PV activation (blue filled circles, magenta dashed line), ChR2-SST activation (blue filled circles, orange dashed line), Arch-PV inactivation (green filled circles, magenta dashed line), Arch-SST inactivation (green filled circles, orange dashed line) and in control (black filled circles, black dashed line) are plotted together for comparison. *In vivo* data represent mean ± SEM. **e** Cosine similarity of iSR-iFR profiles between *in vivo* data in each condition and simulation of the network model with FFI (top), FBI (middle) and with FFI:FBI ratio varied between 0 and 1 to find the optimal ratio for the synchronization of spike-times (bottom). Magenta bar: optogenetic modulation of PV interneuron *in vivo*. Orange bar: optogenetic modulation of SST interneuron *in vivo*. Blue and green shades indicate ChR2 activation with blue light and Arch inactivation with green light in each interneuron subtypes, respectively. **f** Cartoon summarizing the frequency-selective gating of spike-timing synchronization by PV and SST interneurons through preferential activation of FFI and FBI.

## Discussion

Combining single-unit recordings with optogenetic modulation of PV and SST interneurons, we examined the contribution of inhibitory interneurons to the spatio-temporal synchronization of precise spike-times in S1 between L4 and either L5 or L6. We found that on any given trial, some L5 and L6 neurons (which we dubbed “synchronized neurons”) responded to whisker stimulation with spike-timing sequences that replicated both the spike-times and firing-rates of L4 neurons (Fig. 1), showing synchronization across layers. Moreover, we showed for the first time that PV and SST interneurons make different contributions to the synchronization of spike-times from L4 to L5 and L6. PV interneurons helped to synchronize spike-times from L4 to L5 and L6 when the iFR of ISI is low (<12 Hz, Fig. 2k, 3k) whereas SST interneurons help to synchronize spike-times when the iFR of ISI is high (>12 Hz, Fig. 2l, 3l). We then ran *in silico* simulations of three-layer spiking neural networks with different degrees of feedforward and feedback inhibition (Fig. 4). These simulations revealed that the impact of PV and SST manipulations on spatio-temporal spike-timing synchronization *in vivo* mirrored the impact of *in silico* manipulations of feedforward and feedback inhibition, respectively.

One of the most important, novel aspects of our study is the examination of how spike-timing synchronization depends on the iFR of ISI (Fig. 1l, 2k, 2l, 3k, and 3l). If we had limited our analysis to the synchronization of spike-times irrespective of their iFR, we would not have observed any differences between PV and SST interneurons. This result provides further evidence that, *in vivo*, spike-timing and firing-rate cannot be considered in isolation^7–9^. Therefore, we propose that future research into the communication of sensory information using spike-times should incorporate iFR analyses.

By using both optogenetic activation and inactivation of interneurons, we were able to demonstrate that PV and SST interneurons serve as firing-rate-dependent filters on the synchronization of spike-timing sequences between L4 and L5/L6 (Fig. 2k, 2l, 3k, and 3l). Specifically, PV interneurons function as a low-pass filter, helping to synchronize spike-times when firing-rates are low, whereas SST interneurons function as a high-pass filter, helping to synchronize spike-times when firing-rates are high. Given our results and recent findings that the activity of these interneuron classes modulates different oscillatory states^43–45^, future work should examine how these distinct interneuron classes modulate information communication via spike-times synchronization during different oscillatory brain states, e.g. when gamma or theta rhythms are more prominent.

By using *in vivo* spike-timing sequences as input to our *in silico* model and comparing the simulated and real data, we found that our *in vivo* data can be explained if PV interneurons preferentially contribute to feedforward inhibition while SST interneurons preferentially contribute to feedback inhibition, which is surprisingly consistent with *in vitro* and *in vivo* studies of these interneuron classes in S1^21–27^. Moreover, the low/high-pass filtering function of PV and SST interneurons in our study is also consistent with reports that feedforward inhibition filters high-frequency spikes^46,47^ and even vetoes the propagation of high-frequency epileptic activity^48^ while feedback inhibition synchronizes spikes to gamma frequencies^49^. However, considering that only a fraction of neurons are modulated by light^50^ and that SST interneurons can directly inhibit PV interneurons^21,51,52^, it could be that silencing of SST interneurons enhanced the disinhibited PV interneurons (Fig. 3k, l). This complicates any direct assignment of PV interneurons to feedforward circuits and SST interneurons to feedback circuits (Fig. 4), so caution is warranted in these interpretations.

There are several additional limitations to our study that should also be noted. First, differences in photo-addressability, connectivity and distribution of PV and SST interneurons across layers ^21^ and how they are affected under different behavioral/modulatory states may complicate interpretations of their role in spike-timing synchronization^21,26,51,53,54^. While many *in vivo* behavior studies revealed that spike-timing synchronization is related to the behavioral context^1–6,55^, our *in vivo* experiments are conducted under anesthesia. Future studies will need to address our findings in the awake state. Nevertheless, our results from lightly anaesthetized animals may be of significance to neuronal processing and memory consolidation in sleep states. Second, the three-layer network model that we used here does not capture the intricate anatomical and physiological characteristics of the real S1, meaning all conclusions drawn from the model must be taken with some caution. Despite such shortcomings, we used the network model solely to examine the synchronization of spikes across layers, and this is a well-established modeling framework that is widely used in investigating the synchronization of different neural codes^41,42^. Third, based on the canonical microcircuit for the flow of sensory information in S1^30,32,33^, we focused our analyses on the synchronization of spike-times between L4 and the sub-granular layers, L5 and L6. However, many L5 neurons have spike latencies as short as L4 neurons, suggesting that primary thalamic information flows simultaneously to L4 and L5^34^. Therefore, the synchronization of sensory information and spike-times via other routes deserves further investigation.

In summary, by combining single-unit recording, optogenetics and computational modeling, we have shown that temporally precise spike-times can be reliably synchronized from L4 to L5 and L6 during passive whisker stimulation in an anesthetized state *in vivo*. Additionally, we have shown that such synchronization is dynamically gated by PV and SST interneurons, possibly due to their preferential recruitment of distinct inhibitory motifs. Our results add to the repertoire of proposed functions of PV and SST interneurons, and help to delineate their potential role in gating the sensory information processing across cortical laminae.

## Acknowledgement

This work was supported by the Human Frontiers Science Program (HFSP) Young Investigator Award (RGY0073/2015) to B.A.R., M.M.K. & J.K. and by the National Research Foundation of Korea (NRF-2019M3E5D2A01058328) to J.K. H.J.J. was supported by a Korea University Grant. J.M.R. was supported by a Wellcome Trust Prize Studentship.

## Author Contributions

H.J.J., B.A.R., M.M.K. and J.K. conceived and designed the *in vivo* study. H.J.J., M.M.K., H.C. and J.M.R. performed *in vivo* experiments. H.J.J. and H.C. analyzed the *in vivo* data. H.J.J. and J.K. designed the network model and H.J.J. performed the simulations and analysis. J.M.R. and M.M.K. performed immunohistochemistry. H.J.J. and J.K. wrote the original draft. H.J.J., B.A.R., M.M.K., and J.K. reviewed and edited the manuscript.

## Methods

### Animals

All experimental procedures were approved by the Institutional Animal Care and Use Committee (IACUC) at Korea University (KUIACUC-2017-157) and all experimental procedures at the University of Oxford involving animals were conducted in accordance with the UK animals in Scientific Procedures Act (1986). We used three different lines of mice, C57BL/6 wild-type (Gyerim experimental animal resource center in Korea, Envigo in the UK), PV-Cre (JAX#017320, Jackson Laboratory) and SST-Cre (JAX#013044, Jackson Laboratory) mice of either sex. Mice were maintained in a temperature-controlled environment on a 12h/12h light/dark cycle. Food and water were provided *ad libitum*.

### Virus and stereotaxic surgery

To optogenetically modulate the spike activities of PV and SST interneurons (Fig. 2, 3, and Supplementary Fig. 3, 5 and 6)^39^, we expressed blue light-gated cation channel (Channelrhodopsin-2, ChR2) and green-yellow light-gated H^+^ transporter (Archaerhodopsin-3, Arch) in these neuronal types by injecting either adeno-associated virus (rAAV) vectors AAV5-EF1a-DIO-hChR2(E123T/T159C)-mCherry (UNC Vector Core)^39^ or AAV5-EF1a-DIO-eArch3.0-EYFP (UNC Vector Core)^40^ in PV-Cre mice (postnatal days (p) 46-84) and SST-Cre mice (p 49-83). Mice were head-fixed into a stereotaxic device (51730, Stoelting Inc.) under deep isoflurane anesthesia and viral solutions (500-600 nl, ChR2: 3.8×10^12^ virus molecules/ml, Arch: 5×10^12^ virus molecules/mL) were delivered to either the left or right barrel cortex (S1, AP: −/+3.3 mm, ML: 1.3 mm from bregma). Injections were made at two cortical depths (300 and 600 μm depth from pia) using either a 5 μl syringe connected to a motorized stereotaxic injector (The Stoelting Quintessential Injector, 53311, Stoelting Inc.) or a bevelled injection pipette connected to a nanolitre injector (Nanoject II, Drummond Scientific) at a speed of 50-150 nL/min. The injection needle was left in the brain for more than five minutes following injection to prevent withdrawal of virus. At least two weeks of recovery time was allowed after the surgery before conducting *in vivo* recordings.

### *In vivo* recording and optogenetic light-stimulation

Mice were head-fixed into a stereotaxic device (51730, Stoelting Inc.) under anesthesia (ketamine (75-100 mg/kg) and medetomidine (1 mg/kg)). In vivo single-unit recordings were made by implanting a 32-channel silicon probe (A1×32-poly2-5mm-50s-177-OA32, A1×32-5mm-25-177-A32, Neuronexus; Fig. 1a, b and Supplementary Fig. 1) into S1 on either left or right hemisphere (AP: −3.3 mm, ML: 1.3 mm from bregma, 800-950 μm depth from the pia) during stimulation of whiskers on the contralateral side of the electrode implant. Whiskers were glued together and inserted into a capillary tube attached to a piezoelectric bimorph actuator (E650.00 LVPZD amplifier, Physik Instrumente) or motorized actuator which was controlled by custom-made pulse generator based on Arduino and stimulated sinusoidally at 12 Hz for 3 s (Fig. 1a). Recordings of whisker stimulation-evoked responses were repeated 40 times with an inter-trial interval of 10 s. Body temperature was monitored and maintained at 37 °C using a DC temperature control system (40-90-8D, FHC Inc.) throughout all experiments. Signals were sampled at 25 kHz (RZ2 system, Tucker-Davis Technologies) or 30 kHz (RHD2000, Intan Technologies).

Blue light (473 nm) diode laser (iBeam-smart-473, Toptica Photonics) was used to activate ChR2-expressing PV (ChR2-PV) and SST (ChR2-SST) interneurons^39^ and green light (565 nm) LED (M565F3 with LEDD1B, Thorlabs) was used to inactivate Arch-expressing PV (Arch-PV) and SST (Arch-SST) interneurons^40^ (Fig. 2, 3, and Supplementary Fig. 3, 5 and 6). Light was delivered through an optical fiber (Diameter: 200 μm, 0.22 NA, FG200UCC, Thorlabs) placed on the cortical surface <500 μm from the recording site or through an optic fiber which was laminated on the 32-channel optrodes (A1×32-poly2-5mm-50s-177-OA32, Neuronexus). To prevent light stimulation-evoked artifacts, light stimulation preceded the onset of whisker stimulation by 500 ms (Fig. 2e, 2f, 3e, and 3f). Optogenetic activation of ChR2-PV and ChR2-SST interneurons could shut down spike activities of excitatory neurons while optogenetic inactivation of Arch-PV and Arch-SST interneuron could induce epileptic spike activity in excitatory neurons^56^. In order to determine the light stimulation intensity for the modulation of PV and SST interneurons, we measured the firing rates of excitatory neurons during whisker stimulation combined with light stimulation of blue laser (499.7 mW/mm^2^, measured at fiber tip, Supplementary Fig. 5a-d) and green LED (166.2 mW/mm^2^, measured at fiber tip, Supplementary Fig. 5e-h). We used light stimulation intensities of 5, 10, and 50% of the maximal power of the blue laser and at 25-75% of the maximal power of the green LED since these intensities modulated the firing rates of excitatory neurons with little variability (Standard deviation in PV-Cre mice with blue laser = 5.93%, SST-Cre mice with blue laser = 8.05%, PV-Cre mice with green LED = 4.13%, SST-Cre mice with green LED = 4.64%) without excessive suppression or activation of excitatory activities (Supplementary Fig. 5). The effect of blue and green light stimulation to ChR2- and Arch-PV and SST interneurons in sub-granular layers (L5 and L6), respectively, were confirmed by their firing rate changes upon light stimulation (Supplementary Fig. 6).

### Immunohistochemistry

Following *in vivo* recordings, mice were decapitated and their brains were fixed overnight in 4% paraformaldehyde in phosphate buffered saline (PBS, Sigma Aldrich). Brains were then washed in PBS before being cryoprotected overnight in 10% sucrose in PBS. Tissues were frozen in dry ice and 50 μm coronal cryosections were made around the injection site in S1. The fluorescent tags of the ChR2 and Arch opsins, mCherry and EYFP, respectively, were boosted by counter-staining for the tag. Slices were washed with PBS and incubated for 2 h in blocking solution (0.1 M PBS, 0.25% Triton, 5% normal goat serum, Sigma Aldrich). Tissue was treated with primary antibody, rabbit anti-dsRed (Takara Bio) 1:500 or chicken anti-GFP (Abcam) 1:1000, overnight at 4 °C in blocking solution. After washing in PBS, slices were stained with secondary antibody, goat anti-rabbit Alexa Fluor 568 (Life Technology) or goat anti-chicken Alexa Fluor 488 (Abcam), 1:1000 for 2 h at room temperature. Tissue was washed in PBS before counterstaining with 4’,6-diamidino-2-phenylindole (DAPI, Thermo Fisher Scientific) and mounting, with the slides sealed using nail polish. Images were acquired using confocal microscopy (LSM880, Zeiss, Fig. 1b, 2b, and 3b).

### Analysis of *in vivo* electrophysiology data

All data were analyzed offline using custom written codes in MATLAB (R2017a). To determine the laminar location of each single-unit (Fig. 1b), we estimated the cortical depth of the individual channels using the current source density (CSD) depth profiles from local field potentials (LFPs, Supplementary Fig. 1)^28^. LFPs were analyzed by down-sampling the raw signals to 1 kHz and band-pass filtering them at 0.5-300 Hz. We averaged LFPs recorded during whisker stimulations^28^ and applied spatial smoothing and secondary derivation on the averaged LFP by following equations to analyze CSD depth profile:

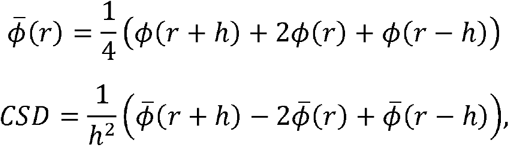

where *ϕ*(*r*) is LFP at depth *r* and *h* is the depth interval. Positive value corresponds to sources and negative value corresponds to sinks in the CDS depth profiles (Supplementary Fig. 1). L4 was identified as early sink in upper layer and L5B was identified as early sink in lower layer^28^ (Supplementary Fig. 1).

To obtain single-unit spike activities from the *in vivo*-recorded raw signals (Fig. 2c), raw signals were band-pass filtered (300-5,000 Hz) after which spike detection and sorting were performed using Klusta-suite software^57^. The spike threshold level was set to 4.5-folds of the standard deviation of each recorded signal. To ensure the single-unit isolation quality, we inspected the shape of spike waveforms, inter-spike intervals, and the shape of the auto-correlogram of spike-times^58^. We only included units showing clear negative deflection in the spike waveform, from which spike waveforms −0.5 to +1.0 ms relative to the spike peak were extracted. Also, units with inter-spike intervals not violating the refractory period of neurons (2 ms) were considered as a single unit^28,58^. We calculated auto-correlogram of spikes using the following equation:

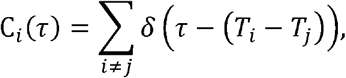

where *T_i,j_* is the spike-times *i* and *j* and we used units having clear refractory periods.

We classified the neuronal types into putative excitatory neurons and inhibitory interneurons by calculating the initial baseline-to-peak amplitude (*a*, Fig. 1c), the last baseline-to-peak amplitude (*b*, Fig. 1c), the spike width (*c*, Fig. 1c), and the asymmetry index ([(*b* − *a*)/(*b* + *a*)]) from the spike waveform (Fig. 1c). Neurons that were located on the right side of the decision boundary ([(*b* − *a*)/(*b* + *a*)] = 2 * *c* − 0.6, Fig. 1c) were considered putative excitatory neurons while those on the left side of the decision boundary were considered putative interneurons. These putative excitatory neurons were further classified into whisker stimulation-responsive excitatory neurons when their firing rate during whisker stimulation increased more than 2-folds of standard deviation of firing rate 1 s prior to whisker stimulation (Fig. 1d). Peri-stimulus time histogram (PSTH) was computed by averaging spike-times relative to whisker stimulation for each trial using 100-ms time bin. Only putative excitatory neurons that were classified as whisker stimulation-responsive were used for further analysis.

### Analysis of the synchronization of precise spike-times

To identify the first cortical layer that responds to whisker stimulation, we analyzed the peak latency of whisker stimulation-evoked multi-unit activity (MUA) in each layer. MUA was defined as the median spike-times of all neurons in each layer within a 50-ms time window after whisker stimulation onset^29^ (Fig. 1e). Although there was no statistical difference between the peak MUA latencies in each layer, following the canonical flow of sensory information, we assumed L4 as input to S1.

To determine whether precise spike-times and spike-timing sequences of units in L4 can synchronize to sub-granular layers (L5 and L6 of S1), we calculated how similar spike-times of L4 and that of other neurons in sub-granular layers (Fig. 1f) and supra-granular layers (L2/3, Supplementary Fig. 4) are using two different measures: spike-time coherence score and similarity (SR). Spike-time coherence score measures the degree of correlation between binarized spike-timing sequences (Fig. 1g, h). We performed pair-wise coherence analysis of L4 neuron and a neuron in another layer through taking the Fourier transform of the cross-covariance function of binarized (2-ms bin) spike-timing sequence (*s_x_*(*t*) and *s_y_*(*t*)) and normalizing it by the Fourier transforms of the auto-covariance function as follow:

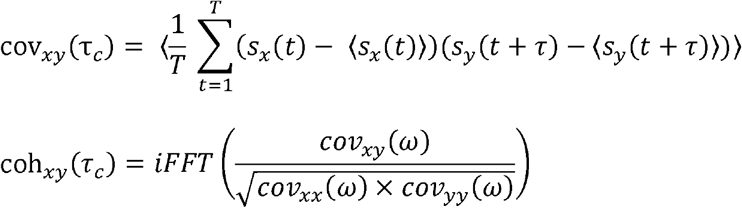

where < > is expected values, *T* is the total duration of *s_x_*(*t*), *τ_c_* is time lag and *cov*(*ω*) is Fourier transform of *cov*(*τ*)^59^. To reliably find precise synchronization of spike-times evoked by whisker stimulation across the layers within cortical column, peak coherence within a narrow time window (−10 ms <*τ_c_*<10 ms) that showing whisker stimulation-evoked columnar responses ^28^ was used as coherence value for further analysis. After calculating the spike-time coherence score of each pair of neurons, we analyzed the probability distribution of spike-time coherence scores and its multimodality was tested by Silverman’s unimodal test^60^ (Fig. 1h, 2g, 2h, 3g, and 3h). If the distribution of spike-time coherence was not statistically unimodal, we fit a mixture of two log-normal distributions to spike-time coherence distribution using the Expectation-Maximization algorithm^36^. The intersection of the two log-normal distributions was defined as the empirically determined “threshold” (Fig. 1h, 2g, 2h, 3g, and 3h) that helped us classify the neurons into two groups, “synchronized neuron” and “non-synchronized neuron” (Fig. 1i). The statistical differences between the distributions of spike-time coherence scores between layers and experiment conditions (Fig. 1h, 2g, 2h, 3g, and 3h) were compared using two-sample Kolmogorov-Smirnov test^61^. The coherence analysis was performed in 1 s-long time-window that slid in 10 ms-steps across the 3 s-long whisker stimulation-evoked spike-timing sequence and the 1 s-long spike-timing sequences showing maximum coherence was used for further analysis.

In order to investigate whether there is a trial-to-trial variability in the role of a given neuron in synchronized or non-synchronized spike-timing synchronization, we counted the number of switches made over trials of a non-synchronized neuron to synchronized neuron or vice versa (Supplementary Fig. 3a-c). We calculated the probability of switch as dividing the number of switches by to the total number of trials (Supplementary Fig. 3d).

To further analyze whether the precise spike-times of L4 could synchronize to “synchronized neurons” in the sub-granular layers in a given trial, we used a measure called SR, which quantifies the synchronization spike-times of two spike trains^37^. SR was calculated by normalizing the number of quasi-simultaneous spike-times between the two spike-timing sequences using the following equations:

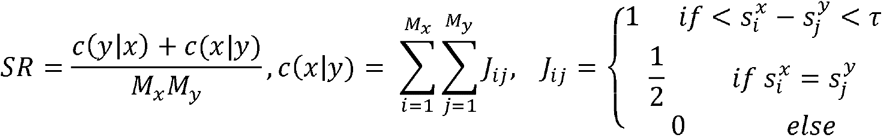

where *τ* is the maximal synaptic time delay (20 ms), *M_x,y_* is the number of spikes for spike-timing sequence *x* and *y*, 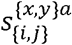 is spike-times of *i* or *j*th spike-timing sequence *x* or *y*. SR ranged from 0 to 1 where 0 means two spike-timing sequences have no simultaneous spike-times and 1 means two spike-times are identical within a synchronization timing window (**±**10 ms) which is an upper limit of within-column activity^4,31^. In order to capture the transient, instantaneously-changing characteristics of the spike-timing sequence, we introduced a new measure we termed instantaneous similarity (iSR) which quantifies SR within the inter-spike intervals of two consecutive spike-timing sequences of L4 (Fig. 1k) as follows:

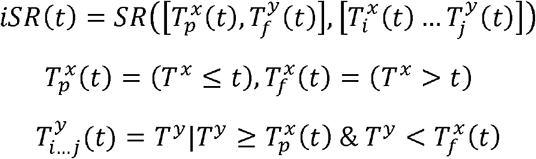

where *SR*(*x, y*) is SR calculation for spike-timing sequence *x* and *y*, *T^x,y^* is spike-times of *x* or *y*, 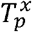 is the previous spike at time *t*, 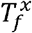 is the following spike at time *t* of spike-timing sequence *x*, and 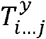 is the subset spike-times of spike-timing sequence of *y* within 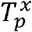 and 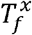.

The transient, instantaneous firing-rates (*iFR*(*t*)) was analyzed by taking the reciprocal of inter-spike intervals. *iFR*(*t*) was categorized into 4 distinct frequency ranges (0.5-4 Hz, 4-12 Hz, 12-20 Hz, 20-50 Hz, Fig. 1j, 2i–2j, and 3i–3j). To quantitatively characterize the frequency-dependent synchronization of whisker stimulation-evoked spike-timing sequence, we plotted *iSR*(*t*) as a function of *iFR*(*t*) which we termed ‘iSR-iFR profile’ (Fig. 1l, 2k, 2l, 3k, and 3l).

To test if the synchronization of precise spike-times we observed with the whisker stimulation-evoked spikes is a biological phenomenon that occurs above chance level, we generated two surrogate data sets by shuffling the inter-spike intervals of the whisker stimulation-evoked spikes recorded *in vivo* (Supplementary Fig. 2a-c)^62^ and generating spike-timing sequences through Poisson-distributed random process (Supplementary Fig. 2d-f). For shuffling the inter-spike intervals, we generated a random sequence of inter-spike intervals by keeping the first spike-time and the number of spikes. For the Poisson-distributed random process, we generated random spike-times while keeping the number of spikes the same.

### Simulation of computational network model

We computationally modeled a three-layer network model composed of excitatory and inhibitory neurons. Each layer was composed of two hundred excitatory neurons and fifty inhibitory interneurons (Fig. 4a), reflecting the fact that interneurons comprise 20-25% of cortical neurons^63^. Excitatory neurons and inhibitory interneurons were modeled as a single compartment conductance-based Hodgkin-Huxley neuron model^64^. Membrane potential of excitatory neuron (*V_m,EX_*(*t*) and inhibitory interneuron (*V_m,IN_*(*t*)) are described by the following equations:

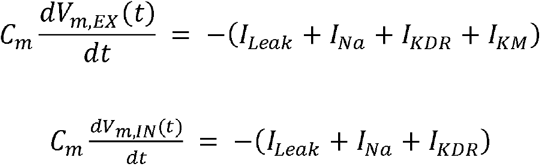

where *C_m_* is the membrane capacitance, *I_Leak_* is the leak, *I_Na_* is the fast sodium channel current^65^, *I_KDR_* is the delayed-rectifier potassium channel^65^, and *I_KM_* is the M-type potassium channel current^66^. The parameters used for excitatory neurons and inhibitory interneurons are shown in Table 1.

**Table 1.**
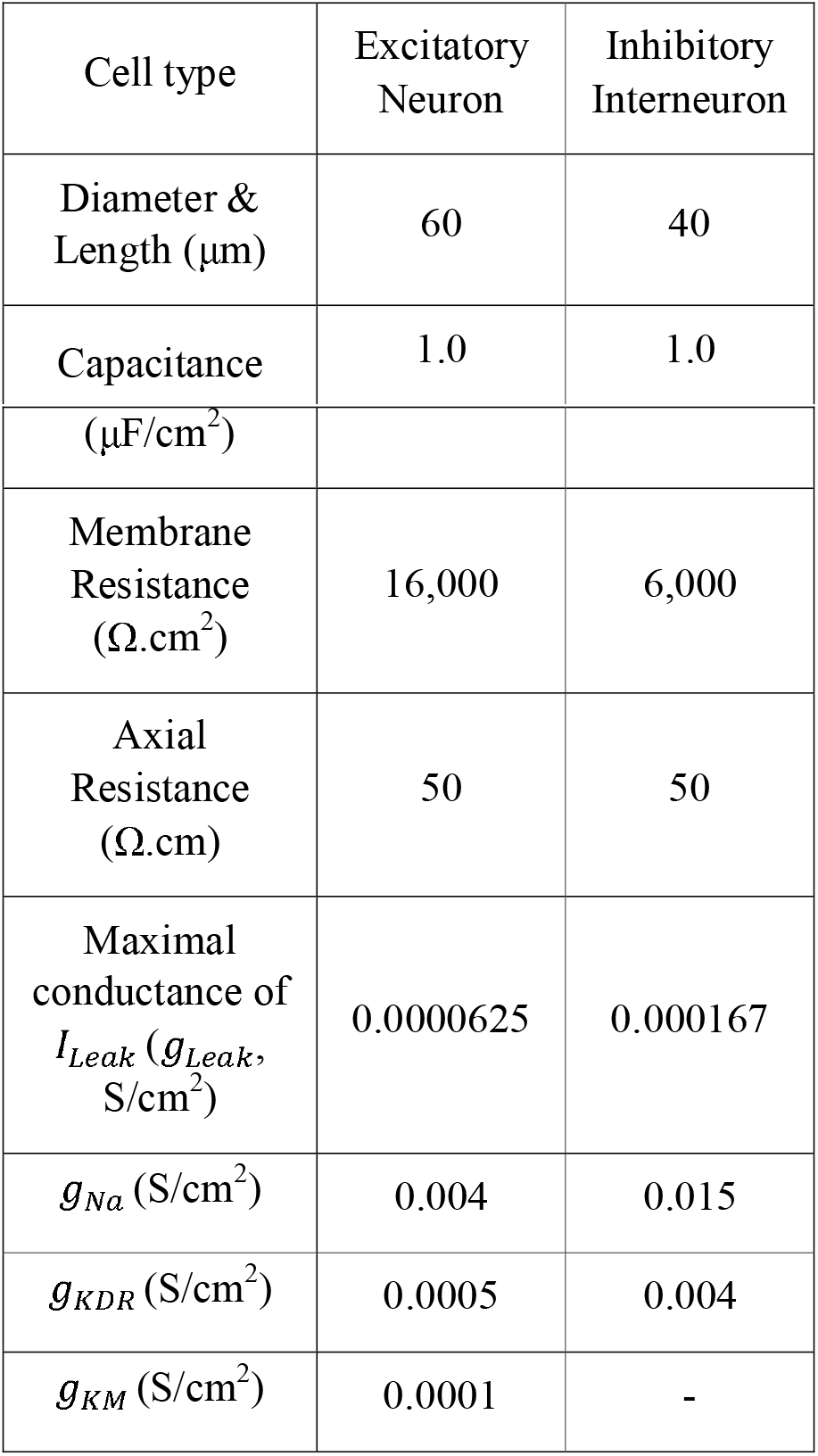
Membrane properties in the Hodgkin-Huxley type excitatory neuron and inhibitory interneuron model.

| Cell type | Excitatory Neuron | Inhibitory Interneuron |
| --- | --- | --- |
| Diameter & Length ($\mu\text{m}$) | 60 | 40 |
| Capacitance | 1.0 | 1.0 |
| ($\mu\text{F}/\text{cm}^2$) | | |
| Membrane Resistance ($\Omega.\text{cm}^2$) | 16,000 | 6,000 |
| Axial Resistance ($\Omega.\text{cm}$) | 50 | 50 |
| Maximal conductance of $I_{Leak}$ ($g_{Leak}$, $\text{S}/\text{cm}^2$) | 0.0000625 | 0.000167 |
| $g_{Na}$ ($\text{S}/\text{cm}^2$) | 0.004 | 0.015 |
| $g_{KDR}$ ($\text{S}/\text{cm}^2$) | 0.0005 | 0.004 |
| $g_{KM}$ ($\text{S}/\text{cm}^2$) | 0.0001 | - |

Each excitatory neuron within a layer was modeled to receive excitatory synaptic inputs from randomly-chosen excitatory neurons in the previous layer with connectivity probability of 10% (Fig. 4a), reflecting the fact that cortical neurons have 10% connectivity^67^. Each excitatory neuron was modeled to receive equal number of excitatory inputs and inhibitory inputs to balance excitation and inhibition of the network, similar to *in vivo* observations from cortical neural network^16^. For the excitatory and inhibitory synapses, double-exponential conductance model was used as following equation:

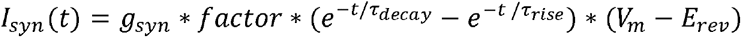

where *g_syn_* is the maximal conductance of synapse, *factor* is the normalizing constant, *τ_decay_* is the decay time constant, *τ_rise_* is the rise time constant, *V_m_* is the membrane potential, and *E_rev_* is the reversal potential of the synapse model. Each parameter was chosen to satisfy the unitary postsynaptic potentials observed *in vitro*^68^ (Table 2).

**Table 2.** Synapse model properties between excitatory neurons and inhibitory interneurons.

| Synapses<br>(presynaptic cell-<br>postsynaptic cell) | Excitatory neuron-Excitatory neuron | Excitatory neuron-Inhibitory neuron | Inhibitory neuron-Excitatory neuron |
| --- | --- | --- | --- |
| Maximal<br>conductance<br>($g_{syn}$, nS) | 0.0074 | 0.0012 | 0.00743 |
| Rise time constant<br>($\tau_{rise}$, ms) | 0.1 | 0.2 | 1.0 |
| Decay time constant<br>($\tau_{decay}$, ms) | 0.2 | 0.3 | 4.5 |
| Reversal potential of<br>synapse ($E_{rev}$, mV) | 0 | 0 | -70 |

To investigate the role of inhibitory circuit motifs in the synchronization of precise spike-times, three different inhibitory network structures were designed: a three-layer network model with inhibitory interneurons providing (1) feedforward inhibition (Fig. 4a, top), (2) feedback inhibition (Fig. 4a, middle), or (3) both feedforward and feedback inhibition at different ratios (Fig. 4a, bottom). Feedforward inhibition was modeled such that interneurons received shared afferent excitatory inputs from excitatory neurons from the previous layer. Feedback inhibition was modeled such that interneuron received excitatory inputs from the excitatory neurons within the same layer, which in turn recurrently inhibited the excitatory neurons.

We modeled sparse representation of sensory information in cortical neurons ^35^ by giving *in vivo* spike-timing sequence recorded from L4 during whisker stimulation (Fig. 1) as input to only forty excitatory neurons in the input layer (L_input_) of the network model while the remaining one hundred and sixty excitatory neurons were modeled to spontaneously spike with Poisson-randomized spike-times with log-normally distributed firing rates (μ= 3.5, σ = 2.5) as observed *in vivo* (Fig. 4b)^69^.

To infer which inhibitory circuit motif is mediated by PV and SST interneurons, we treated iSR-iFR profile as 4D vector and compared 4D vector of *in vivo* data and simulation data by calculating cosine similarity (Fig. 4e) using the following equation:

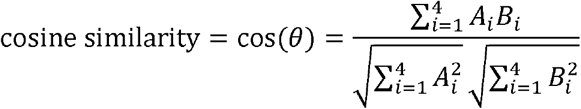

All simulations were conducted with sampling rate of 5 kHz in NEURON simulation environment^70^.

### Statistical Analysis

All values were represented as mean ± SEM, where ‘n’ refers to the number of animals from which *in vivo* recording were made or the number of simulations ran for computational simulation. For determining statistical significance of iSR-iFR profiles between synchronized and non-synchronized neuron, we used Wilcoxon rank sum test showing the significance of iSR at each iFR frequency (Fig. 1l, 2k, 3l, 3k, and 3l). The statistical difference of iSR between iFR bands, we used one-way ANOVA test (Fig. 1l). For the comparison of coherence distribution, we used Kolmogorov-Smirnov test. The resulting two-tailed *p* value that was less than 0.05 was deemed statistically significant.

## Data Availability

Klusta-suite is an open source spike detection and sorting software downloadable at https://github.com/kwikteam/klusta. Custom-made MATLAB codes used for data analysis are available from the corresponding authors upon request.

