## Supplementary Figures for "Distinct roles of parvalbumin and somatostatin interneurons in the synchronization of spike-times in the neocortex"

**Supplementary Figures 1 - 6**

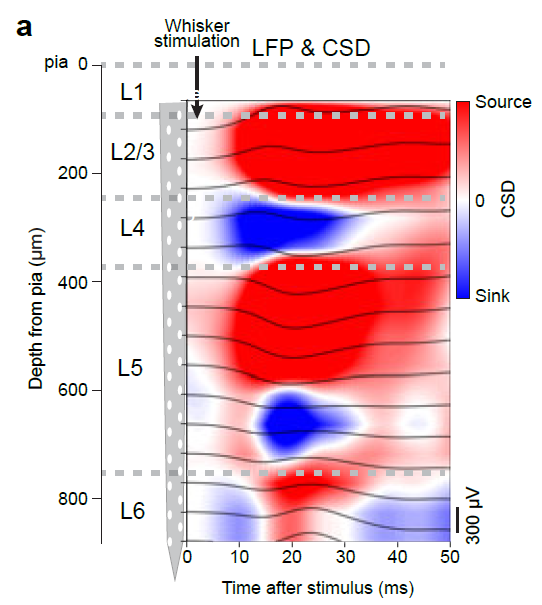

**Supplementary Figure 1.** Current-source density analysis to estimate the depth of 32-channel silicon probe in S1. **a** Schematic of a 32-channel silicon probe with an example of local field potential (LFP, black solid) and current-source density (CSD) plot during whisker stimulation (red represents source and blue represents sink).

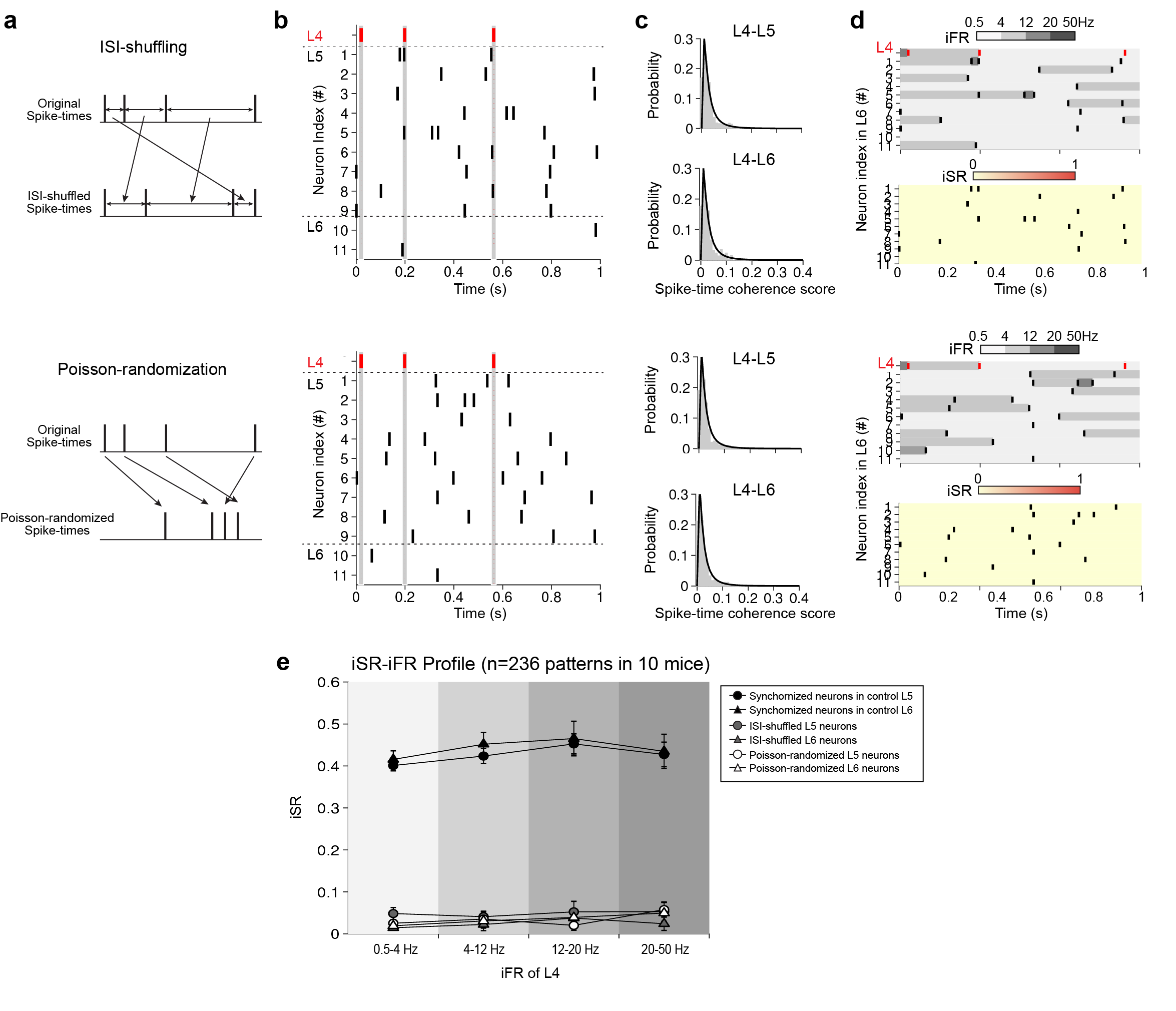
**Supplementary Figure 2.** ISI-shuffled and Poisson-randomized spike-times fail to synchronize. **a** Schematic diagrams illustrating inter-spike intervals (ISI)-shuffling (top) and Poisson-randomization (bottom) of spike-times. **b** Raster plots showing spike-timing patterns of each neuron in L4, L5 and L6 as recorded *in vivo* as in Figure 1F (left), after ISI-shuffling (top) and Poisson-randomization (bottom) of spike timing-patterns in L5 and L6. Gray shade indicates the synchronization time window (±10 ms). **c** Distribution of pair-wise spike-time coherence scores of after ISI-shuffling (top) and after Poisson-randomization (bottom) of spike-timing patterns in pairs of L4-L5 and L4-L6. **d** Example of instantaneous firing rate (iFR) and instantaneous similarity (iSR) between spike-timing patterns of L4 and L6 after ISI-shuffling (top) and after Poisson-randomization of spike-timing patterns (bottom). iFR is classified into four different frequency bands (0.5 - 4Hz, 4 - 12 Hz, 12 - 20 Hz and 20 - 50 Hz) presented in grayscale. iSR differences are represented by the red color-scale (bottom). **e** iSR-iFR profile for spike-timing patterns of L4-L5 (circle) and L4-L6 (triangle) pair for control (black), after ISI-shuffling (gray) and after Poisson-randomization (empty) of spike-timing patterns.

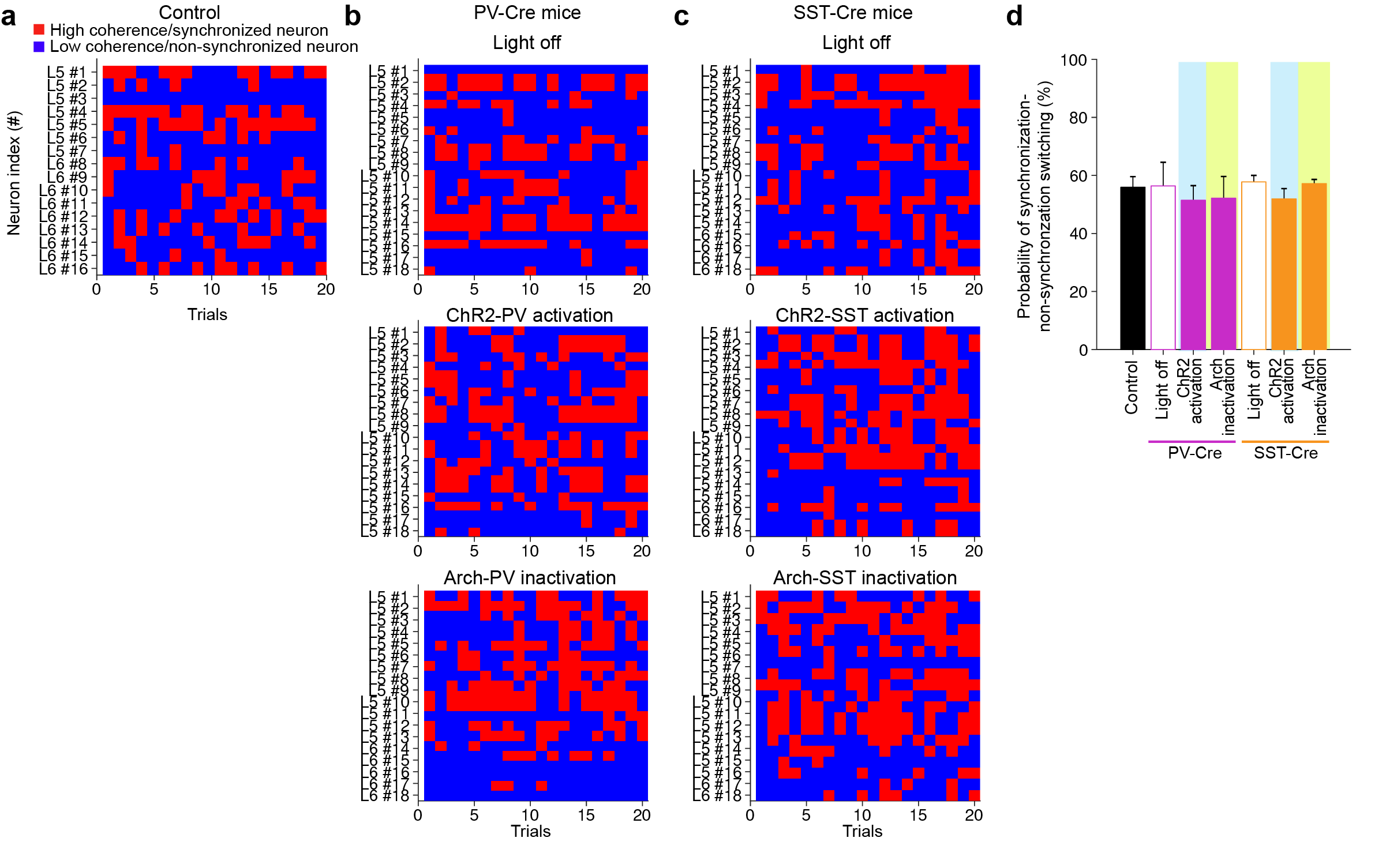
**Supplementary Figure 3.** Trial-to-trial variability of spike-time coherence score. **a** 2-D color map representing each neuron alternating in its role as a synchronized neuron (red) or a non-synchronized neuron (blue) over trials during whisker stimulation in the control mice. **b, c** Same as (**a**) but in PV-Cre mice (**b**) and in SST-Cre mice (**c**) without (top) and with the activation of ChR2-PV (**b**, middle), the inactivation of Arch-PV (**b**, bottom), the activation of ChR2-SST interneurons (**c**, middle), or inactivation of Arch-SST (**c**, bottom). **d** The probability of switching between synchronized and non-synchronized neuron over trials in each condition. Data are mean ± SEM.

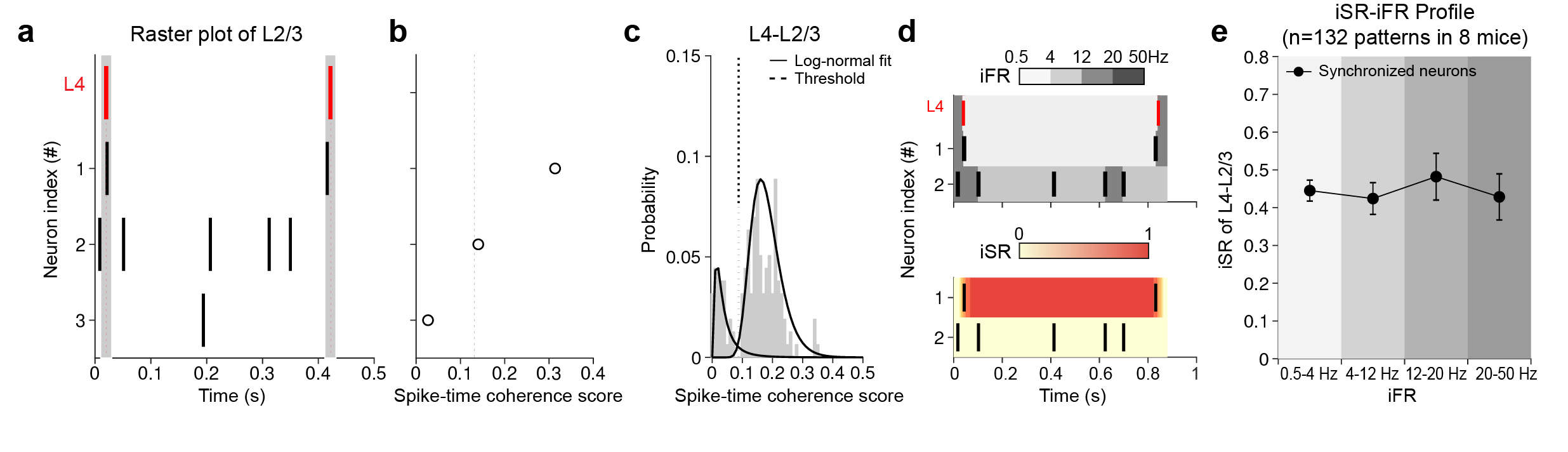
**Supplementary Figure 4.** Synchronization of whisker stimulation-evoked spikes from granular to supra-granular layers in S1. **a** Raster plots of neurons in L2/3 during whisker stimulation. Gray shade indicates the synchronization time window (±10 ms). **b** Spike-time coherence scores of each pair of L4-L2/3 spike-timing patterns. Vertical dotted line represents the empirically defined threshold decision boundary between synchronized and non-synchronized neurons (see Supplementary Fig. 4c). **c** Distribution of spike-time coherence scores over trials for each pair of L4-L2/3 spike-timing patterns. Threshold: intersection between two log-normal distributions (dotted line). **d** Examples of instantaneous firing-rate (iFR, top) and instantaneous similarity (iSR, bottom) between spike-timing patterns of L4 and L2/3. iFR is classified into four different frequency bands (0.5 - 4Hz, 4 - 12 Hz, 12 - 20 Hz and 20 - 50 Hz) presented in different gray colors. iSR is represented in red color-scale (bottom). **e** iSR plotted as a function of iFR in iSR-iFR profile for spike-timing patterns of L4-L2/3. All data are mean ± SEM and n represent number of animals.

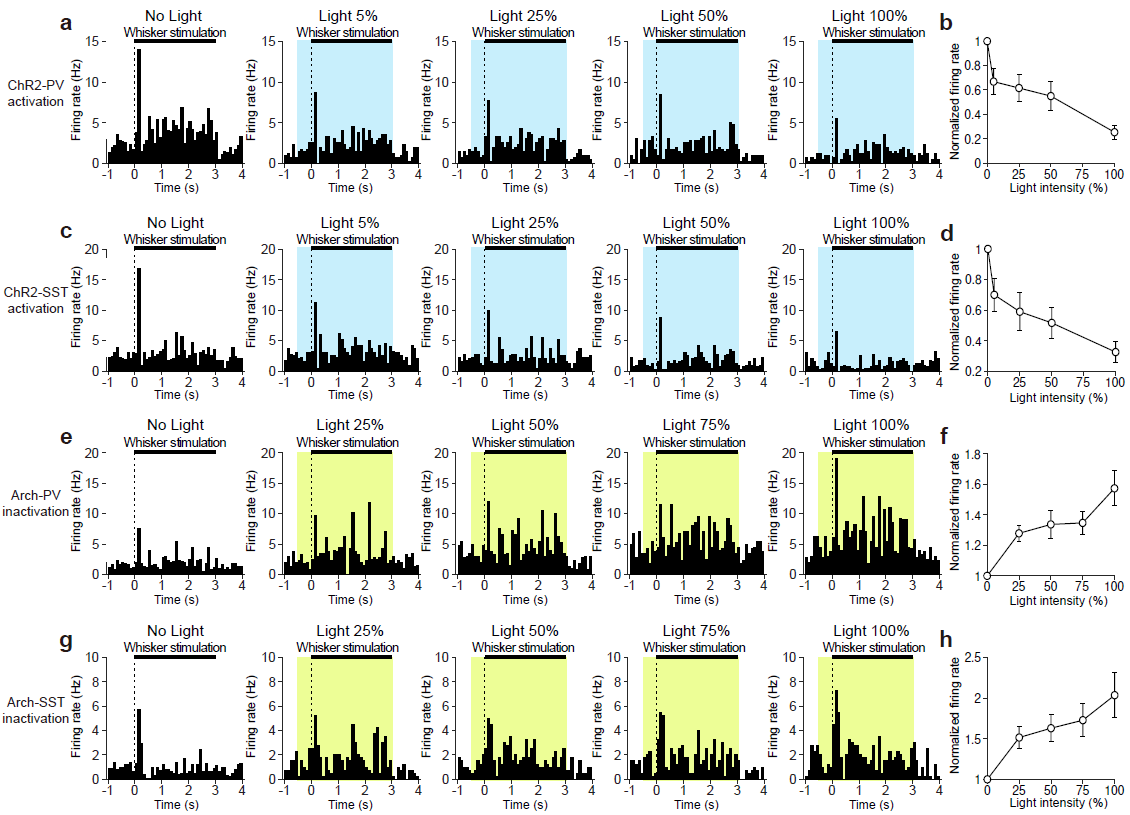

**Supplementary Figure 5.** Light stimulation intensity-dependent firing rate changes in ChR2 and Arch-expressing PV and SST interneurons. **a** PSTHs of putative excitatory neurons (black vertical bars) during whisker stimulation (black horizontal bar) in response to blue light stimulation (473 nm, blue shade) of different laser intensities (in % of the maximal power 499.7 mW/mm^2^) to activate ChR2-PV interneurons in PV-Cre mice. **b** Spike firing rates normalized to firing rate without light stimulation and plotted as a function of laser light intensity in PV-Cre mice. **c, d** Same as (**a, b**) but with ChR2-SST interneurons in SST-Cre mice. **e** PSTHs of putative excitatory neurons (black vertical bars) during whisker stimulation (black horizontal bar) in response to green light stimulation (565 nm, green shade) of different LED intensities (in % of the maximal power 166.2 mW/mm2) to inactivate Arch-PV interneurons in PV-Cre mice. **f** Spike firing rate normalized to firing rate without light stimulation and plotted as a function of LED light intensity in PV-Cre mice. **g, h** Same as (**e, f**) but in Arch-SST interneurons in SST-Cre mice.

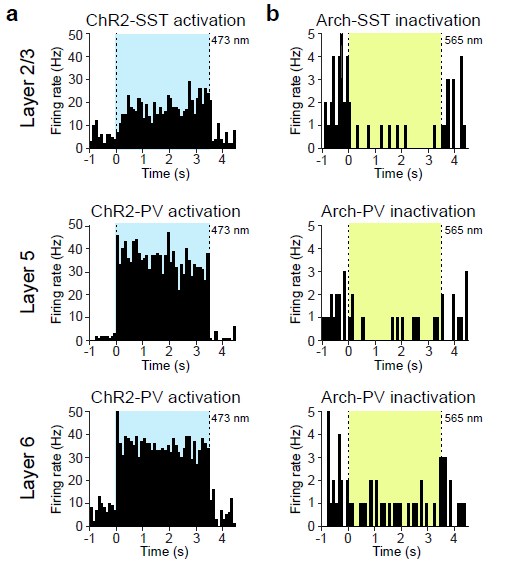

**Supplementary Figure 6.** Optogenetic activation and inactivation of PV and SST interneurons across layers in S1. **a** Example PSTHs of ChR2-PVs and ChR2-SST interneurons showing increase in firing rate in response to blue light (473 nm, blue shade) in L2/3, L5 and L6.

**b** Example PSTHs of Arch-PVs and Arch-SST interneurons showing decrease in firing rate in response to green light (565 nm, green shade) in L2/3, L5 and L6.
